# Genomic and transcriptomic analysis of sacred fig (*Ficus religiosa*)

**DOI:** 10.1101/2022.06.21.497063

**Authors:** K. L. Ashalatha, Kallare P Arunkumar, Malali Gowda

## Abstract

**Background:** Peepal/Bodhi tree (*Ficus religiosa* L.) is an important, long-lived keystone ecological species. Communities on the Indian subcontinent have extensively employed the plant in Ayurveda, traditional medicine, and spiritual practices. The Peepal tree is often thought to produce oxygen both during the day and at night by Indian folks. The goal of our research was to produce molecular resources using whole-genome and transcriptome sequencing techniques.

**Results:** The complete genome of the Peepal tree was sequenced using two next-generation sequencers Illumina HiSeq1000 and MGISEQ-2000. We assembled the draft genome of 406 Mb, using a hybrid assembly workflow. The genome annotation resulted in 35,093 protein-coding genes; 53% of its genome consists of repetitive sequences. To understand the physiological pathways in leaf tissues, we analyzed photosynthetically distinct conditions: bright sunny days and nights. The RNA-seq analysis supported the expression of 26,479 unigenes. The leaf transcriptomic analysis of the diurnal and nocturnal periods revealed the expression of the significant number of genes involved in the carbon-fixation pathway.

**Conclusions:** This study presents a draft hybrid genome assembly for *F. religiosa* and its functional annotated genes. The genomic and transcriptomic data-derived pathways have been analyzed for future studies on the Peepal tree.

## Background

The Peepal tree (*Ficus religiosa* L.) is a sacred fig, hemi-epiphyte that belongs to the Moraceae family and has a diploid sporophytic chromosome count (2n = 26) [1]. It is known to be a long-lived deciduous species related to the 755 fig species widespread worldwide [2]. The Peepal tree is a cosmopolitan species, having value for cultural and spiritual practices in Buddhism, Hinduism, and Jainism. It is popularly called the Bodhi tree, where Buddha is believed to have meditated and attained spiritual enlightenment underneath this tree. Hence, the culture is spread across Asia and it has been worshipped. Peepal has several vernacular names, like Pippali, Ashwatha, Arali, and so on; it is frequently found together with the Neem tree near Indian temples [3]. Generally, the Peepal tree has a special significance in communities across India as it is believed to produce oxygen day and night. They have a special type of stomata called sunken, giant, or hydathode at the lower leaf epidermis. These are larger than the normal stomata and occur over the veins or are mixed with normal stomata. It indicates that such stomata hold gaseous and water molecules for a longer time [4]. To our knowledge, there is no reported scientific evidence to claim oxygen production from the Peepal tree at night.

In Ayurveda, the Peepal tree has been classified as a Rasayana (a type of drug), whereby rejuvenators and antioxidants aid in relieving the body’s stress [5]. Peepal tree alleviates Pitta and Kapha (Ayurvedic classifications), hence prescribed for treatment of the disorders like respiratory and inflammatory disorders, ulcers, stomatitis, hiccup, arthritis, gout, skin diseases, bone fracture, diabetes, etc., [5]. In animal models such as rats, the Peepal tree has been tested for the treatment of neurodegenerative disorders such as Parkinson’s disease and Huntington’s disease [6] [7], as well as anti-ulcer activity in albino mice [8].

Next-generation sequencing (NGS) technologies have accelerated the generation of draft genome sequences of Moraceae plant species. The genome size of *Morus notabilis* is 330 Mb [9], 333 Mb in *Ficus carica* [10], 436 Mb in *F. microcarpa* and 370 Mb in *F. hispida* [11]. The genome sequencing of non-model plant species *F. religiosa* was first mentioned in The Neem Genome book chapter [3]. A recent study has generated the genomic resource of *F. religiosa* (332 Mb) and *F. benghalensis* (392 Mb). However, they generated a limited size of genome assembly and genes (23,929) for the Peepal tree when compared to the present study [12]. Recently, a few research groups have attempted sequencing of non-model plant species like pineapple (*Ananas comosus*) and *Kalanchoë* species revealing the gene expressions of the Crassulacean acid metabolism (CAM) pathway [13] [14]. The whole-genome sequencing has shown common or crystalline ice plants (*Mesembryanthemum crystallinum*) to switch from Calvin-Benson Cycle (C3) to CAM photosynthesis under a salt stress [15]. The study described by comparing both species with and without the C4 trait and different tissues within a C4 plant using RNA-seq suggests ways of integration into the underlying C3 metabolism [16]. These findings and other physiological features of the Peepal tree enabled us to characterize the pathways in the present work.

Despite its ecological, medicinal, cultural, and historic importance, the molecular biology and genomics studies on the Peepal tree are scanty. As Peepal is relevant to traditional medicinal practices and Buddha’s meditation, we envisaged elucidating the genome sequence and studying the transcriptome of photosynthetic tissue (leaf tissue) in diurnal and nocturnal periods. The objective of the present study was to generate a genome sequence and annotate genes of the Peepal tree. The transcriptomic analysis has been undertaken to identify the expression of genes in the diurnal and nocturnal periods for photosynthetic activity using a molecular approach. In this study, we aimed to characterize the genes involved in various physiological, biochemical metabolic, and other pathways. Also, a comparative genomic analysis has been carried out to study the relationship of the Peepal tree with closely related species of its Moraceae family.

## Results

### *De novo* hybrid assembly using Illumina and MGI short reads

We used two next-generation technology platforms to sequence the whole genome of the fig species, the Peepal tree. A total of 266 and 645 million paired-end reads were generated from Illumina HiSeq1000 and MGISEQ-2000 platforms respectively. The data of 88.44 billion high-quality bases (Quality>20) was used for genome assembly. A hybrid assembly was performed using a sequencing depth of 65.5X Illumina reads and 158.86X MGI reads. The raw data details are given in the supplementary material (Additional file 1: Table S1.1). The evaluation of the distribution of k-mers in both Illumina and MGI reads to estimate the genome size provided genome sizes of 319 Mb and 273 Mb respectively (Additional file 2: Figures S1A and S1B). The combination of Illumina and MGI reads was used for assembling the genome. Hybrid genome assembly yielded a genome of 406 Mb. The contig N50 length is 5,817 bp and the largest contig length is 148 Kb. The GC content of the Peepal tree genome is 34.23%. The gap-closing step was performed for the hybrid assembly. There were 35,811 (5.5%) misassembled contigs and 604,807 (94.4%) truly assembled contig sequences in the final assembled genome. The workflow of the genome assembly is presented in the supplementary material (Additional file 3: Figure S2). The statistics of assembly contigs and scaffolds are shown in the supplementary material (Additional file 4: Table S1.2) and the final scaffold assembly of the genome is given in Table 1. The alignment of raw reads to the hybrid genome sequence was performed, which mapped 99.5% and 99.27% of Illumina and MGI reads respectively (Additional file 5: Text file S1.1).

**Table 1:**
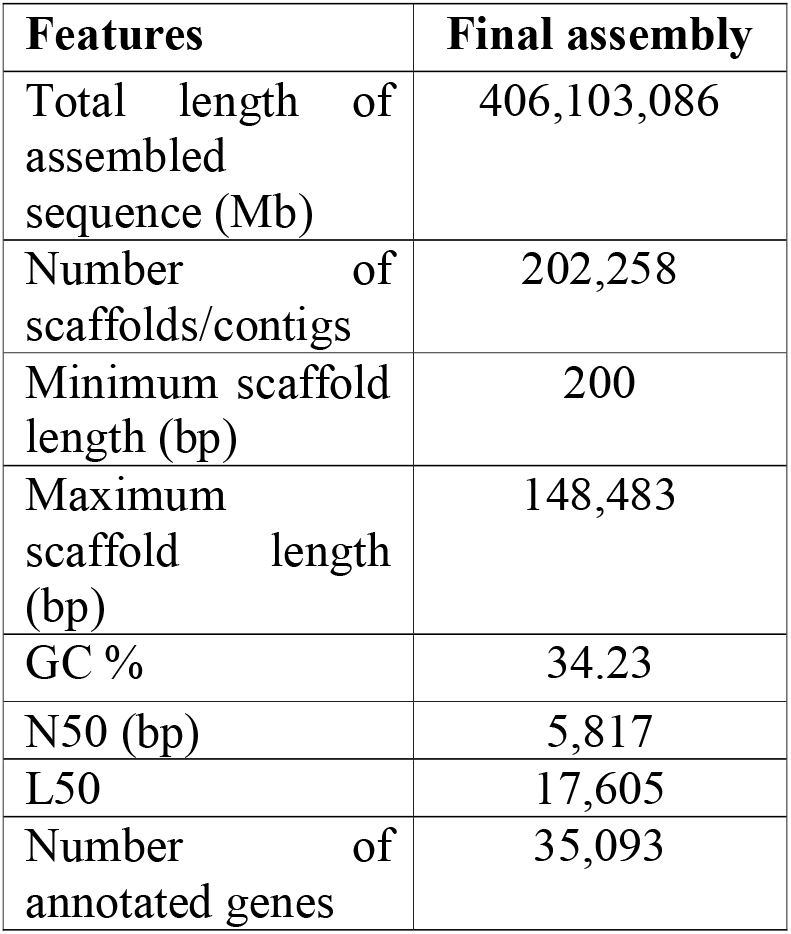
Final assembly and annotation of Peepal genome

The completeness of the Peepal tree genome assembly was assessed with the BUSCO tool. The results showed that 76.5% (232 out of 303) and 84.1% (1,210 out of 1,440) of genes were conserved as single-copy orthologs in eukaryotic and plant universal data sets, respectively. Out of 232 complete genes in the Eukaryota database, 214 are single-copy orthologs, 18 are duplicates, 57 are fragmented and 14 are missing. Out of 1,210 complete genes in the Embryophyte database, 1,173 are single-copy orthologs, 37 are duplicates, 105 are fragmented and 125 are missing (Additional file 6: Figures S3A and S3B). The transcriptome sequence reads aligned with the assembled genome showed that 99.46% of all reads were mapped and of these 88.25% of paired reads were mapped (Additional file 7: Text file S1.2).

### Genome and pathways annotation

We identified 35,093 protein-coding genes with the complete structures in the Peepal tree genome (Additional file 8: Text file S2.1 and Additional file 9: Text file S2.2). RNA-seq data from two leaf tissue samples of the Peepal tree and alternative reference ESTs from *Morus notabilis* and *Arabidopsis thaliana* protein sequences were used as protein homology evidence during genome annotation. Based on a homology search using BLASTN, out of 35,093 genes predicted, 32,255 genes (91.9%) were having evidence from transcriptome assembly. About 76.3% of RNA-seq reads from the day and night leaf tissue samples were mapped to the annotated genes in the Peepal tree.

Based on the sequence similarities, the complete set of annotated genes and their amino acid sequences were used in the Kyoto Encyclopedia of Genes and Genomes (KEGG) pathway analysis [17]. This result showed the pathways in metabolism, biosynthesis of secondary metabolites, genetic and environmental information processing, and signal transduction pathway were common as several others. The top 5 highest gene count for pathways like Ribosome (123 genes), Spliceosome (96 genes), Oxidative phosphorylation (86 genes), Thermogenesis (82 genes), and RNA transport (74 genes) was found. In addition, important candidate genes were also found for human disease pathways like Huntington’s disease (68 genes), Parkinson’s disease (57 genes), Alzheimer’s disease (55 genes), and others (Additional file 10: Table S2).

### Protein family and Gene Ontology analysis

The protein family (Pfam) ID and Gene Ontology (GO terms) were assigned to genes using an InterProScan module [18]. Out of 35,093 genes, 24,163 consisted of Pfam IDs that were distributed across 3,759 types of Pfam domains, and their gene ontology (GO) terms were also identified. The Pfam domain consisting of proteins that were large in the Peepal tree genome included 3-Deoxy-D-manno-octulosonic-acid transferase, Ring finger domain, PPR repeat family, Helix-loop-helix DNA-binding protein, DYW family of nucleic acid deaminases, Lysine methyltransferase, Putative GTPase activating protein for Arf, Ankyrin repeats and others (Additional file 11: Table S3).

Catalase is an antioxidant enzyme known to catalyze H_2_O_2_ into water and oxygen. We identified the gene sequences for the Catalase gene (FRLM_016351-RA) and its isozyme CAT1 Catalase isozyme 1 (FRLM_016350-RA), (FRLM_012250-RA) in the Peepal genome annotation. Two catalase genes were identified in the differential expression of transcriptome data: the KatE gene known as a monofunctional catalase, and the KatG gene known as a catalase-peroxidase [19]. KatE gene also known as CatB, is differentially expressed during the day and night period with the Fragments Per Kilobase of transcript per Million mapped reads (FPKM) values 937.49 and 1786.02 respectively. KatG gene is differentially expressed during the day and night, with the FPKM values 162.03 and 81.53 respectively. The KatE gene has been reported to be involved in physiological pathways such as glyoxylate and dicarboxylate metabolism, tryptophan metabolism, MAPK signaling pathway – plant, FoxO signaling pathway, and serine-pyruvate transaminase pathway. The KatG gene is involved in tryptophan metabolism, tyrosine metabolism, biosynthesis of secondary metabolism, and drug metabolism pathways.

### Identification of homologous, orthologous, and singleton genes

To understand the gene evolution and relationships among *F. religiosa* and other taxa, we performed homologous and orthologous gene detection analysis for the Peepal tree with an additional 5 species. Homologous gene identification and orthologous clustering of the proteomes of six species, including the model organism *A. thaliana, M. notabilis* (closest relative species of Ficus), and other closely related species of the Moraceae family were selected for the analysis. Based on proteome sequence homology analysis, 29,516 homologous genes were found in *Arabidopsis thaliana*, 29,924 in *Morus notabilis*, 29,750 in *Cannabis sativa*, 29,909 in *Prunus persica*, and 29,830 in *Ziziphus jujuba* with respect to *F. religiosa* proteome sequences (35,093). *F. religiosa, A. thaliana, M. notabilis, C. sativa, P. persica*, and *Z. jujuba* form a cluster of 24,310 orthologous genes and are conserved within the species. The number of specific orthologous gene clusters identified was 15,016 in *F. religiosa*, 16,170 in *A. thaliana*, 16,517 in *M. notabilis*, 15,921 in *C. sativa*, 16,655 in *P. persica*, and 17,235 in *Z. jujuba*. A total of 1,184 single-copy gene clusters were found across the six species and the number of specific singletons identified was 10,154 in *F. religiosa*, 4,469 in *A. thaliana*, 2,284 in *M. notabilis*, 1,912 in *C. sativa*, 1,802 in *P. persica*, and 4,209 in *Z. jujuba* (Figure 1).

**Figure 1:**
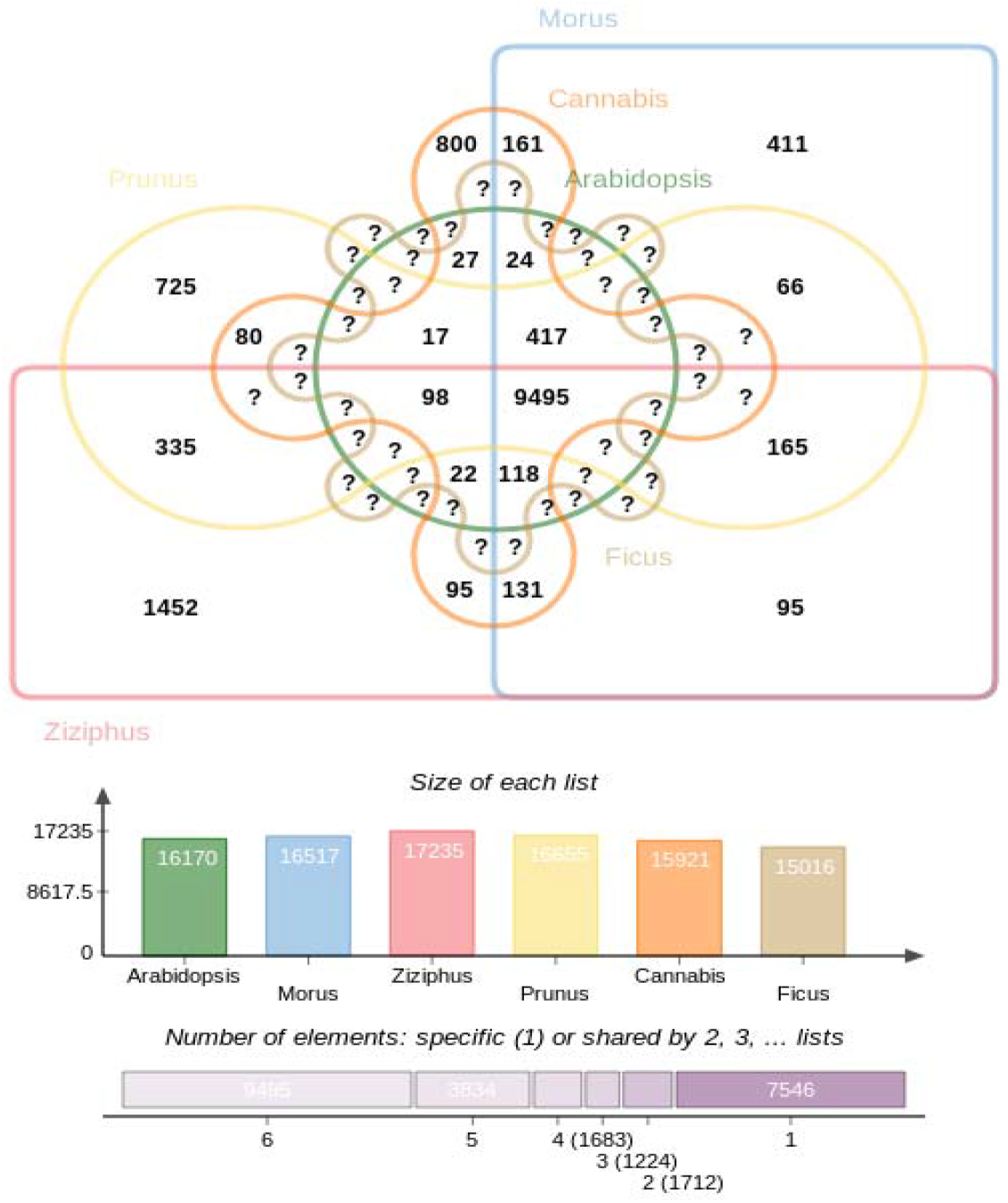
Orthologous clustering of 6 species using proteome data deduced 24,310 orthologous gene clusters and 1,184 single-copy gene clusters across the above 6 species

The identified single-copy clusters were used to illustrate the taxonomic and phylogenetic relationships among a group of species. Based on the similarity of proteomes and single-copy orthologous clustering, we deduced the phylogenetic tree for *F. religiosa* and the other five species. The multiple sequence alignment (MSA) and Neighbour-Joining (NJ) methods were used for constructing an evolutionary phylogenetic tree. It was found that *F. religiosa* is closely related to *M. notabilis* by having more similarities in their proteomes as they are evolving from the Moraceae family, followed by *C. sativa, Z. jujuba, P. persica*, and *A. thaliana* (Additional file 12: Figure S4).

### Comparative analysis of Peepal tree genome

We aligned the genome of the Peepal tree with those of the three Moraceae family members, *F. carica, F. microcarpa*, and *M. notabilis*. Comparison of our assembly against these genomes resulted in a mapping of 88.62% to *F. carica*, 89.6% to *F. microcarpa*, and 46.9% to *M. notabilis*. The results showed Peepal genome to be closer to the genus Ficus (*F. carica* and *F. microcarpa*) and also relatively closer to the genus Morus of the same family. The statistics of genome sequence alignment of *F. religiosa* against *F. carica, M. notabilis, F. microcarpa* genomes is presented in the supplementary material (Additional file 13: Text file S3).

### Repeats in the genome of the Peepal tree

Repeat library building and repeat identification were performed using the ReapeatModeller and RepeatMasker tools (www.repeatmasker.org) respectively. *De novo* repeat identification resulted in 53.55% (269.62 Mb) repetitive sequences in the Peepal tree genome. The RNA elements, long terminal repeats (LTR) constitute about 5% of repeats and 43.71% of these repeats did not belong to any of the annotated repeats families. The 53.55% of repetitive sequences in the Peepal tree genome are closest to its Moraceae family species, 47% are found in the closest species mulberry (*M. notabilis*), 46.5% in *F. microcarpa*, and 48.9% in *F. hispida*. The repetitive sequences were classified into known categories, such as LINE1 (0.19%), long terminal repeat retrotransposon (5.09%), DNA transposons (1.09%), and simple repeats (3.25%) and unclassified (43.71%) (Additional file 14: Table S4).

### Simple sequence repeats (SSRs)

We identified SSRs from the assembled Peepal tree genome. In total, 799,992 SSRs were identified on 267,593 sequences, which are composed of mono- (606,169), di- (143,113), tri- (34,327), tetra- (11,791), penta- (2,911), and hexa- (1,681) type repeats (Additional file 15: Table S5.1). Among mono repeats, the ‘A/T’ (73.91%) type was the highest followed by ‘C/G’ (1.87%). Similarly, the ‘AT /TA’, ‘AG/CT’, ‘AC/GT’, and ‘CG/CG’ types of di repeats were in 9.8%, 2.76%, 1.41%, and 0.09% fractions, respectively. ‘AAT/ATT’, ‘AAG/CTT’ ‘ATA/TAT’, ‘TTA/TTC’, and ‘GAA/TAA’, were the most abundant tri repeats and ‘AAAT’ was predominant in tetra repeats. The detailed distribution of all types of repeats and statistics is shown in the supplementary material (Additional file 16: Table S5.2 and Additional file 17: Table S5.3).

### Transcription Factors (TFs)

Transcription factors act in regulating gene expression driven by several external and internal signals by activating or suppressing the downstream genes. The MAKER annotated protein sequences of Peepal tree genome assembly were used for BLAST analysis with the Plant Transcription Factor Database v5.0 [20] using the *A. thaliana* protein sequence as a reference. A total of 1,264 protein sequences from 35,093 protein-coding genes with genome annotation shows evidence for 56 families of Transcription factors (Additional file 18: Table S6). The TFs families include the ERF, M-type MADS, ARF, DBB, MIKC MADS, WOX, C3H, G2-like, MYB, TALE, B3, HB-other, and MYB-related family proteins. The transcription factors play an important role in regulating growth, developmental processes, and environmental responses in the plant’s [21].

### Transcriptome sequencing, assembly, and annotation

*De novo* transcriptome assembly was performed for the mature leaf samples of the Peepal tree collected during the day and night periods. The assembly was performed for each sample and also a combined assembly was performed for the reads of both samples. The combined transcriptome assembly resulted in 152.8 Mb assembled bases with an N50 length of 2,076 bp and an average transcript length of 1316.83 bp and 42.17% GC content. The transcriptome assembly and annotation workflow are given in the supplementary material (Additional file 19: Figure S5). The statistics of assembly contigs and sequence assembled contigs are also provided in the supplementary material (Additional file 20: Table S7).

The *de novo* assembled transcript sequences (116,038) were processed for annotation. *De novo* assembled transcripts were clustered to exclude the redundant transcripts and identified 26,479 unique transcripts sequences. The statistics of Unigenes are given in Table 2. *De novo* assembled transcripts and unigenes were annotated to find the structural and functional genes. The protein families were identified for the uniquely characterized transcripts of RNA data.

**Table 2:** Statistics of Uni-genes in Peepal Transcript

|  | Sample (Day - 2 PM) | Sample (Night - 2 AM) | Combined Sample |
| --- | --- | --- | --- |
| Number of genes | 22,597 | 23,360 | 26,479 |
| Number of transcripts | 16,912 | 16,780 | 18,173 |
| Percent GC | 46.37 | 46.24 | 46.25 |
| Contig N50 (nts) | 1,404 | 1,407 | 1,374 |
| Median contig length | 813 | 801 | 753 |
| Average Contig (nts) | 1060.06 | 1055.95 | 1017.44 |
| Total assembled bases | 23,954,145 | 24,667,005 | 27,055,237 |

Out of 26,479 transcripts, Pfam IDs for 19,175 were distributed across 3,977 types of protein family (Pfam) domains and their gene ontology (GO) terms were also identified (Additional file 21: Table S8). Pfam IDs and GO terms were assigned to predict the function of unique gene sequences and encoded translated proteins.

The differentially expressed genes (35,182) from the day and night periods of leaf tissue samples of the Peepal tree (Additional file 22: Table S9.1) were used for the pathway analysis. The top 272 highly up-regulated differentially expressed transcripts were identified for diurnal and nocturnal periods (Additional file 23: Table S9.2).

The TFs were identified from the differentially expressed transcripts. From the day sample, 2 transcripts coded for specific TFs like C3H family protein and nuclear factor Y, subunit A7 (NF-YA7), and in the night period sample, the 6 transcripts coded for specific TFs like ERF family protein, CONSTANS-like 2, MYB-related family protein (Additional file 24: Table S9.3). In plants, the nuclear factor-YA has a role in drought stress responses. In rice, NF-YA7 is involved in the drought tolerance pathway which is independent of the Abscisic acid manner [22]. Expression of C3H and NF during the day could influence plant growth and development in the Peepal tree. The Ethylene response factor ERF105 showed as the cold-regulated transcription factor gene of Arabidopsis [23]. In the Peepal tree, ERF, MYB - related family proteins like REVEILLE 1 (RVE1) [24] and late elongated hypocotyl gene (LHY) are expressed during the night period. RVE1 functions in the circadian clock and auxin pathways and LHY maintains the circadian rhythm in Arabidopsis [25]. Both the RVE1 and LHY are found expressed in night-specific Peepal tree transcripts indicating the active circadian rhythms and pathways during the dark time.

### Non-coding RNA genes in the Peepal tree genome

Based on a coding potential calculator (CPC), *de novo-*based assembled transcripts (26,479) were further categorized into protein-coding (19,911) and non-coding (6,568). Based on BLASTN analysis, out of 6,568 non-coding transcripts, 4,219 transcripts targeted genome-annotated genes and 2,349 remained non-coding. A total of 30,973 Cufflinks assembled transcripts (reference-based alignment with genome assembly) were further categorized into protein-coding (7,163) and non-coding (23,810). Out of 23,810 non-coding transcripts, 14,605 were having alignment to genome-annotated genes using BLASTN. Further, categorization of specific day and night sample transcripts resulted in 6,628 (day) and 7,339 (night) protein-coding and 20,528 and 25,494 non-coding transcripts respectively. From these non-coding transcripts, 18,893 (day) and 22,232 (night) transcripts were aligned to MAKER-P predicted genes using BLASTN. The remaining transcripts were considered to be non-coding transcripts, as we did not find any match to predicted gene evidence to support them. Hence, the majority of RNA sequences are found to have protein-coding sequences, while the non-coding genes have been shown biologically relevant in recent years, and deepen studies are needed to understand their functions.

#### miRNAs

microRNAs are a major class of non-coding RNAs. Based on the homology search, we identified the microRNA precursors using the miRbase database (http://www.mirbase.org). These microRNAs belong to MIR396, MIR2916, MIR156, MIR164, MIR6236, MIR166, MIR168, and MIR395 families. Among the identified miRNAs, MIR408 was found to be specific to the night period transcripts of the Peepal tree. MIR 408 was identified on the genes like TPK5 Two-pore potassium channel 5, prfA peptide chain release factor 1, and also on proteins of unknown function in the Peepal genome. MIR408 is a highly conserved microRNA in plants and is involved in enhancing photosynthesis by mitigating the efficiency of irradiation utilization and the capacity for carbon dioxide fixation [26].

The unigene transcripts were used to identify the microRNAs. MIR168 and MIR166 homologs were identified on the two transcripts. We identified the miRNAs on genomic scaffolds based on mapping the transcriptome data to the genome. This provides information on miRNAs specific to the day and night leaf tissue transcriptome. We identified 23 and 25 pre-miRNA expressions in the day and night period respectively (Additional file 25: Table S10.1). The statistics of transfer RNAs (tRNA) were identified in the genome and their details are given in the supplementary material (Additional file 26: Table S10.2).

### Elucidation of carbon fixation pathway in Peepal tree

The study was conducted to analyze the gene expression patterns in the leaf tissues of the Peepal tree under the diurnal (2 PM) and the nocturnal period (2 AM). Through the pathway analysis, the candidate genes for carbon fixation pathways like the CAM pathway, Calvin-Benson cycle (C3) pathway, and C4 - Dicarboxylic pathway were identified and estimated based on their transcript abundance. The transcriptome data contained 20 putative genes involved in the carbon fixation module of CAM, C3, and C4 including the key genes fructose-bisphosphate aldolase class I, fructose-1,6-bisphosphate, phosphoenolpyruvate carboxylase (PEPC/PPC), phosphoenolpyruvate carboxylase kinase (PPCK), NAD+ and NADP+, malate dehydrogenase (MDH) and pyruvate orthophosphate dikinase (PPDK) genes (Additional file 27: Table S11). Gene mapping was completed for CAM and C4 cycle pathways and could not find a mapping for three genes in the C3 cycle pathway. Those three genes, fructose-6-phosphate phosphoketolase (EC 4.1.2.22) and phosphoketolase (EC 4.1.2.9) were purified in *Acetobacter xylinum* [27] and sedoheptulokinase (EC 2.7.1.14) was shown in *Bacillus* species [28]. These three genes were not found in the Peepal tree for the C3 cycle.

The differentially expressed genes from transcriptomic data were mapped to the reference carbon fixation in photosynthetic organisms pathway on the KEGG database (Additional file 28: Figure S6) [29], [30], and [31]. The diagrammatic representation of the genes involved in the carbon fixation pathway is shown in the supplementary material (Additional file 29: Figure S7).

The important genes expressed in the C3 cycle are rubisco and glyceraldehyde-3-phosphate dehydrogenase. Ribulose-bisphosphate carboxylase (RuBP carboxylase or *rubisco*) small chain enzyme that is enriched in leaf tissue collected during the day (2 PM). Rubisco is the most abundant protein in chloroplasts. The glyceraldehyde-3-phosphate dehydrogenase (NADP+) is enriched in day sample leaf tissue, the enzyme responsible for the reversible conversion of glyceraldehyde 3-phosphate to ribulose bisphosphate using ATP, the acceptor for CO2. The trancriptomic genes have been mapped to the C3 cycle except for the three genes mentioned above. [32].

The signature genes responsible for the CAM cycle were expressed in the Peepal tree during the night. The phosphoenolpyruvate carboxylase kinase (PPCK), NAD(P)-ME (maeB), and Malate dehydrogenase (MDH) transcripts were highly enriched in the photosynthetic leaf tissue collected during the night period than the day. It indicates that the Peepal tree adapts to the CAM pathway and can fix nocturnal carbon dioxide using the PEP carboxylase (PEPC) enzyme and accumulate malate by the enzyme malate dehydrogenase. The transcriptomic genes of the Peepal tree have been completely mapped to the KEGG pathway of the CAM cycle.

In the C4 Dicarboxylic cycle, the high expression of glutamate-glyoxylate aminotransferase enzyme (GGAT) in the leaf tissue collected during the night period (2 AM) indicates the photorespiration in the Peepal tree. The carbon fixation begins in the mesophyll cells, where CO2 is converted into bicarbonate. It adds the 3-carbon acid phosphoenolpyruvate (PEP) by an enzyme called phosphoenolpyruvate carboxylase. The product of this reaction is the four-carbon acid oxaloacetate, which is reduced to malate another four-carbon acid [33]. The second highest expression is NADP-malate dehydrogenase (MDH), which converts the oxaloacetate generated by PEPC to malate. The differentially expressed genes from the Peepal tree had a complete mapping to the C4 cycle. The gene expression pattern of the carbon-fixation pathway in the Peepal tree suggests that the plant switches between the C3, C4, and CAM cycles during the diurnal and nocturnal periods. The FPKM and Trimmed Mean of M-values (TMM) values of the differentially expressed genes for the carbon fixation pathway are shown in Figures 2A, B, C, and 3A, B, and C.

**Figure 2:**
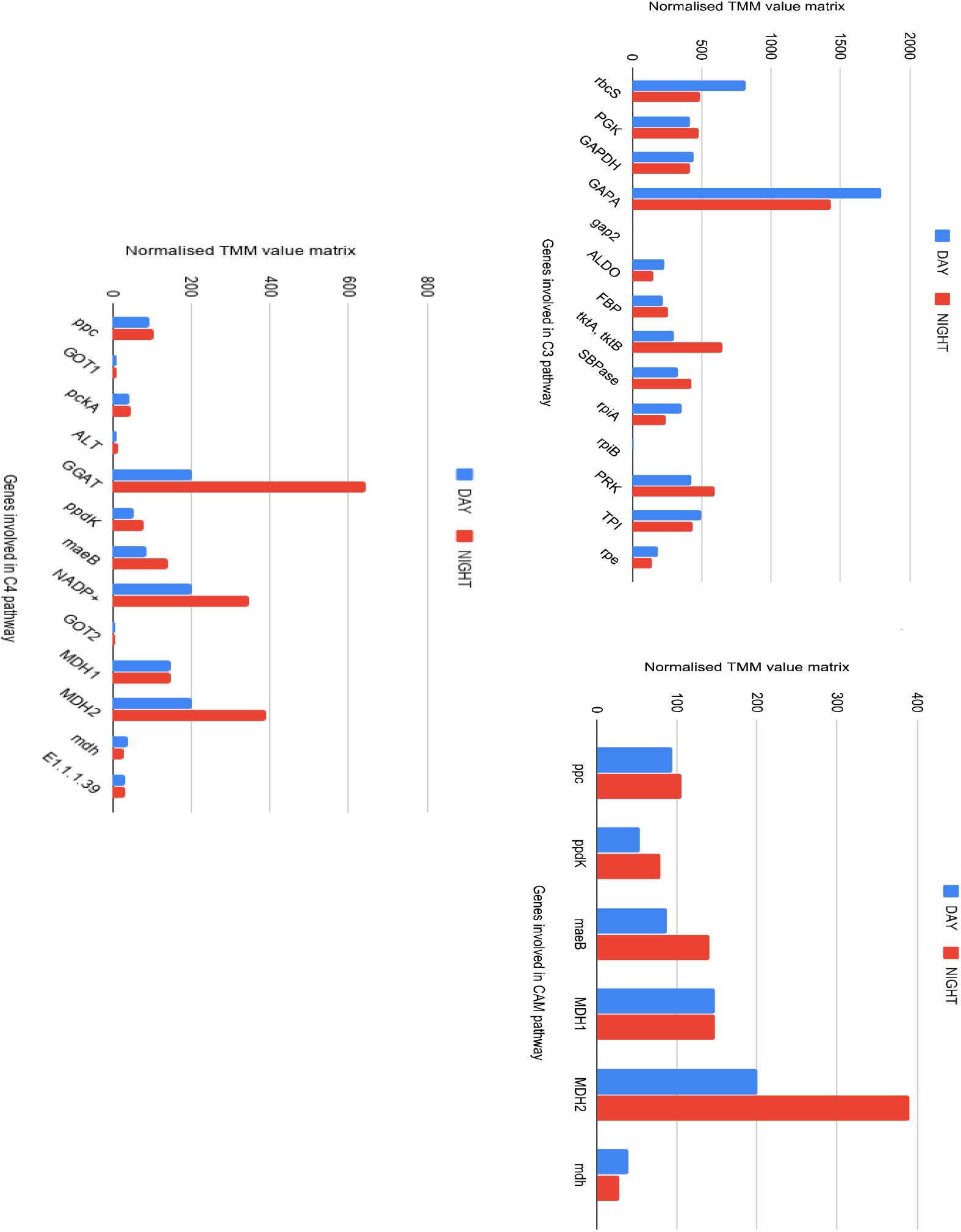
The candidate genes involved in the C3, CAM, and C4 cycles. A) Calvin-Benson (C3) cycle B) Crassulacean acid metabolism (CAM) cycle C) C4 cycle. The X-axis represents the genes involved in pathways, Y-axis is the matrix of normalized expression trimmed mean of M (TMM) values; the Blue graph - leaf tissue collected during the day (2 PM), Red graph - leaf tissue collected during the night period (2 AM).

**Figure 3:**
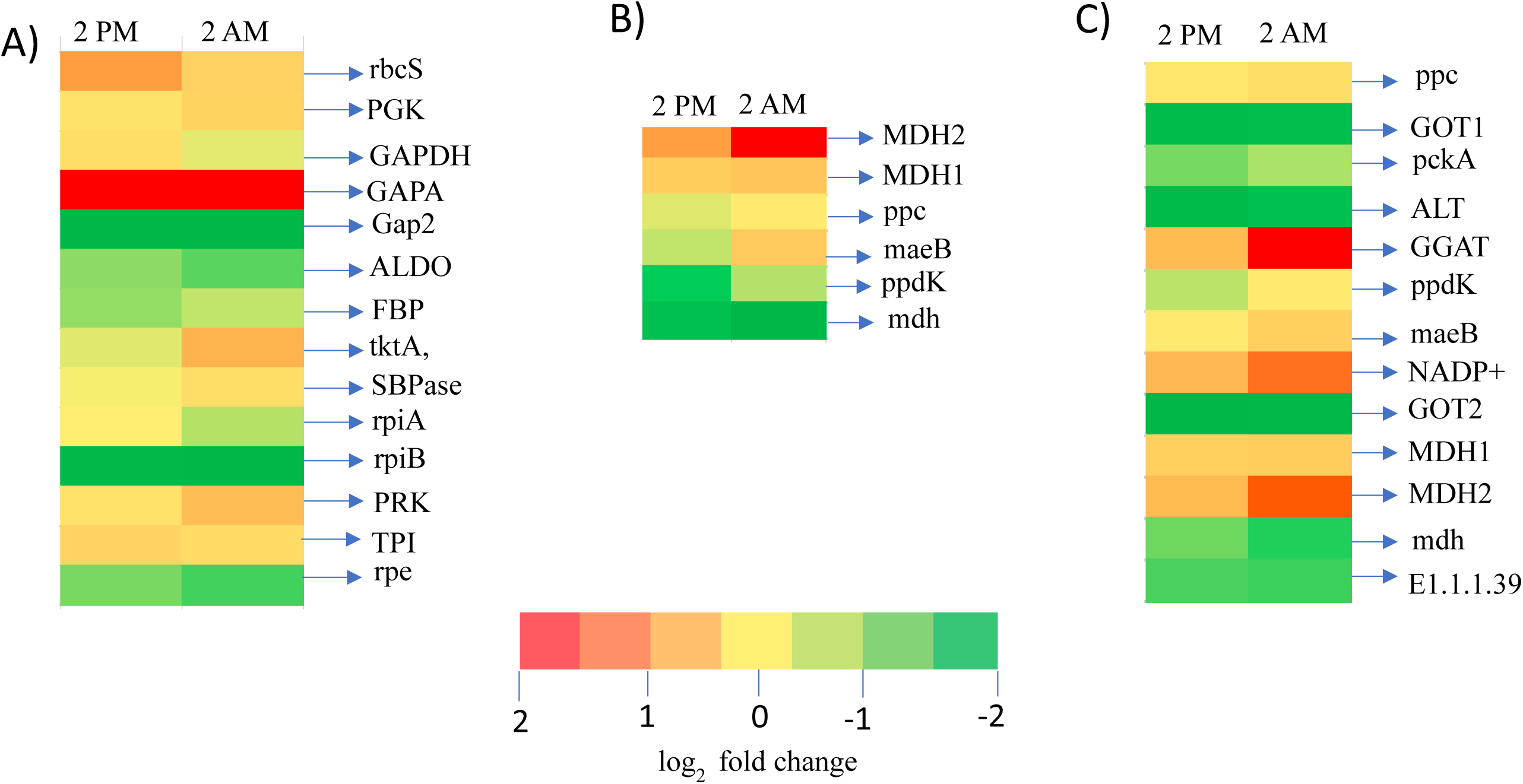
Gene expression pattern of *F. religiosa* carbon fixation genes across the diurnal (2 PM) and nocturnal (2 AM) expression data. A) C3 cycle, B) CAM cycle C) C4 cycle with fold change log2-transformed Fragments Per Kilobase of transcript per Million mapped reads (FPKM) value based expression profiles are shown

## Discussion

This study generated and annotated the genomics and transcriptomics data for the keystone species Peepal tree (*F. religiosa*). We used two next-generation sequencing technologies to sequence the Peepal genome and characterized the hybrid whole genome. The assembled genome resulted in a size of 406 Mb with 35,093 protein-coding genes, based on *ab initio*, homology, and mRNA evidence used for annotation. Photosynthetic tissues at distinct conditions (diurnal and nocturnal) were used for RNA sequencing to understand the genes, proteins, and molecular pathways. The combined transcriptome analysis yielded 26,479 unique transcripts. The completeness of the Peepal tree genome was confirmed based on BUSCO analysis and comparative analysis of transcriptome data.

We performed the downstream analysis of genomic and transcriptomic data to understand the microRNAs, TFs, and molecular pathways of the Peepal tree. The miRNA MIR408 was identified to be specially expressed in the in Peepal leaf tissue during the night period. The miRNA MIR408 responds to copper deficiency and light in Arabidopsis [34]. In *O. sativa*, MiR408 plants were efficient at saving and converting light energy into sugars, suggesting that miR408 can promote photosynthesis by down-regulating the uclacyanin (UCL8) gene [35]. Thus, MIR408 found specific expression in transcripts of night period leaf tissue of the Peepal tree indicating the similar conversion of light energy and accumulation of sugars at night. It may also aid in photosynthesis by enhancing carbon fixation.

In the current study, Catalase gene expression was found to be high in the night period transcripts of Peepal tree leaf tissue. A previous study on the Peepal tree showed that leaf tissue collected at night time exhibited the scotoactive opening of stomata during the night, which indicates that through the stomatal opening molecular oxygen (O_2_) is released by the action of catalase enzyme on hydrogen peroxide (H_2_O_2_) [36]. The physiological interaction between catalase and its substrate H_2_O_2_ in the plant was determined by quantifying H_2_O_2_ and assaying the catalase, in which catalase showed a 4-fold increase in activity, especially during the night. Peepal tree has a higher amount of H_2_O_2_ deposition during the night than day [36], which is an indication of pathway switching between carbon fixation pathways.

The RNA sequencing from diurnal (2 PM) and nocturnal (2 AM) leaf samples showed the gene expression patterns of the carbon fixation pathway. The day mRNA expression data suggested Peepal tree can carry out the diurnal carbon fixation by the C3 cycle. GGAT is involved in the photorespiratory process. High expression of GGAT in the C4 cycle indicates that there could be photorespiration in the Peepal tree during the night. Plants adapt to the CAM cycle to grow during water constraints and increase the level of carbon dioxide uptake than their C3 and C4 cycles [37]. The Peepal tree study provides information on plants using the CAM pathway to fix nocturnal carbon dioxide using the PEP carboxylase (PEPC) enzyme and the accumulation of malate by the enzyme malate dehydrogenase.

The Peepal tree gene expression analysis for the C3, C4, and CAM cycles suggested that plants could switch between these three cycles depending on the carbohydrate, amino acids biosynthesis, metabolism, and environmental conditions. In the *Kalanchoë fedtschenkoi* genome study, the convergence in protein sequence and re-scheduling of diel transcript expression of genes was reported to be involved in nocturnal CO_2_ fixation, stomatal movement, heat tolerance, circadian clock, and carbohydrate metabolism with the other CAM species in comparison with non-CAM species [14]. Some of the previous studies in the pineapple genome revealed the gene lineage transition from C3 photosynthesis to CAM, and CAM-related genes exhibit a diel expression pattern in photosynthetic tissues [38]. The evolution of CAM in *Agave* from C3 photosynthesis shows that the core metabolic components required for CAM have ancient genomic origins which could be traceable to non-vascular plants while regulatory proteins required for diel re-programming of metabolism have shared among the recent origin of C3, C4, and CAM species [39].

The plant model Arabidopsis encodes several orthologues of human proteins that function in mechanisms similar to those in other eukaryotes [40] [41]. Previous findings showed that 70% of oncogenes involved in cancer have orthologs in Arabidopsis, 67% in *D. melanogaster*, 72% in *C. elegans*, and 41% in *S. cerevisiae*. This ‘disease gene’ similarity is comparable to that observed in other model organisms [42]. The research in Arabidopsis and many other model systems has led to the discovery or analysis of genes and processes important to human health. Previous studies showed that the Peepal tree has been tested for the treatment of neurodegenerative disorders such as Parkinson’s disease and Huntington’s disease [6], [7], as well as anti-ulcer activity in animal models. [8]. Another study showed that methanol extract of *F. religiosa* has proven anti-inflammatory properties in LPS-induced activation of BV2 microglial cells, and it might have therapeutic potential for various neurodegenerative diseases [43]. In the present work, we found that Peepal genes show similarities with human disease pathways, which can be utilized to further understand the traditional medicinal practices and Buddha’s meditation practices. Thus, plant research opens up new frontiers in terms of drug development and treatment of diseases of great importance to human health. Plants seem to be a part of this diverse portfolio of tools necessary to understand fundamental cellular processes.

In summary, the genome data and transcript abundance evidence indicate the molecular switch in the carbon fixation pathway of the Peepal tree (*F. religiosa*) during the day and night periods depending on its physiological and environmental conditions. Our study is a foundation for further experiments to determine the underlying mechanisms in C3, C4, and CAM metabolism.

## Conclusions

In this study, we generated the genomic and transcriptomic data for Peepal/Bodhi tree. Genomic data pathway analyses identified the genes associated with several physiological, biochemical, metabolic, and disease pathways. Differential expression data from diurnal and nocturnal leaf tissue samples of Peepal revealed gene expression patterns in the carbon fixation pathway during light and dark. The transcript abundance indicates that plants could switch between the three C3, C4, and CAM pathways. The well-annotated genome for the Peepal tree will have broader implications for studies regarding the physiology, evolution, conservation of species, and human neurological diseases.

## Methods

### Collection of leaf samples and extraction of nucleic acids

The mature leaves were collected from a cultivated Peepal tree (15 years old) at a Private property, Anuganalu village, Hassan District, India (13.0647° N, 76.0363° E). We have followed a non-invasive method for collecting leaf samples. Genomic DNA was extracted from the leaves using the Qiagen DNeasy Plant Mini kit (Catalog #69106), and the quality and quantity of DNA were confirmed using the Nanodrop. From the same Peepal tree, the leaf samples were collected and immediately placed on dry ice during the day (2 PM) and night (2 AM) periods. Total RNA was isolated from the leaf samples using the QiagenRNeasy Plant Mini kit (Catalog #74904) method and was treated with RNase-free DNase I (Catalog #M0303S) from New England BioLabs for 30 min at 37 °C to remove residual DNA. RNA integrity and quantity were confirmed on Qubit and Tape station using a dsDNA HS (Catalog #32854) kit from Invitrogen and RNA screen tape from Agilent respectively.

### DNA and RNA library preparation and sequencing

Whole-genome shotgun DNA library preparation was performed using the Illumina TrueSeq DNA sample preparation kit (FC-121-2001). The paired-end (PE) (2 × 100 nts) sequencing was carried out using Illumina HiSeq-1000. Also, to increase the size of genome data, we sequenced the genome with paired-end (PE) (2 × 100 nts) using the MGISEQ-2000 platform.

The RNA libraries were prepared using “TruSeq RNA Library Prep Kit v2 from Illumina®” with Illumina standardized protocol. The RNA libraries were quantified on Qubit (dsDNA HS kit) and validated on the TapeStation instrument (D1000 screen tape). These RNA libraries were used for sequencing with the Illumina HiSeq-2500 platform.

### Genome size estimation and assembly

Each of the Illumina and MGISEQ-2000 raw reads were processed for a quality check using the FastQC v0.11.6 tool [8]. Then filtering and trimming of raw reads were done to remove the low complexity bases using the TrimGalore-0.4.5 (https://www.bioinformatics.babraham.ac.uk/projects/trimgalore/) and reads having quality value Q>20 and length above 20 bases were taken for constructing the assembly. To estimate the genome size, filtered reads were taken for the k-mer distribution (different k-mers from 21 to 77) and abundance analysis using Jellyfish v1.1.12 [44] and GenomeScope v2.0 [45]. The separate Illumina and MGI Seq generated raw reads were used to construct the assembly using the tools SPAdes-3.13.0 [46] and MaSuRCA-3.2.9 [47] respectively. The parameters were the default k-mer sizes of 21, 33, and 55 for Illumina assembly. The constructed assemblies were used to build the super scaffolds using the tool SSPACE standard v3.0 [48].

The combined Illumina HiSeq and MGISEQ raw reads were used to construct the hybrid assembly using the assembler SPAdes-v3.13.0 [46]. The parameters were the default k-mer sizes 21, 33, and 55, with a 77 mer also set. The gaps in the assembly were closed by GMcloser-1.6.2 [49]. The assembly statistics were obtained using the tool Quast v4.6.1 [50]. The completeness and evaluation of the assembly were done by BUSCOv3 tool [51] with the Embryophyte and Eukaryota database and by aligning the RNA-seq reads to the genome.

### Structural gene prediction and functional annotation

Peepal tree assembled scaffolds were processed for structural and functional gene annotation using the MAKER-P v.2.31.10 software [52]. The RNA-sequenced data of *Morus notabilis* [9] consists of expression sequence tags (ESTs) and the GFF (Gene finding format) file which contains the gene features and structures of genes, protein data of *A. thaliana* and RNA-sequence data of Peepal tree were imported as evidence for annotation support. The structural and functional annotation of predicted genes and proteins was performed using BLASTP in the Uniprot database. The protein family, structures, and gene ontology (GO) terms were identified for protein-coding genes using InterProScan-V5.27-66.0 [18].

### Gene family construction, identification of homologous and orthologous genes

Protein sequences of *A. thaliana, M. notabilis, P. persica, C. sativa, Z. jujuba*, and the protein sequences of the current study *F. religiosa* were taken for the homologous and orthologous gene identification. The homologous genes were identified in *F. religiosa* proteome sequences using the BLASTP analysis against the other 5 proteomes of *A. thaliana, M. notabilis, P. persica, C. sativa*, and *Z. jujuba*. OrthoVenn2 [53] was used to cluster orthologous genes and identify the single-copy orthologous genes in all six proteomes. Further, these single-copy orthologous genes were used for constructing the phylogenetic tree using the tool MAFFT-v7 [54].

### Comparative genome analysis

We downloaded the genomes of *F. carica, F. microcarpa*, and *M. notabilis*. We aligned these genomes against the Peepal tree genome assembly to understand their relationships using the BWA-V0.7.17 (Burrows-Wheeler Aligner) [55] and Samtools v1.7 [56].

### Prediction of repetitive elements: TEs and SSR

The RepeatModeller-open-1.0.11and RepeatMasker-4.0 tools were used for repeat library building and repeat identification in the assembly respectively. The MicroSAtellite identification tool (MISA) [57] was used for the identification of SSRs from assembled genome sequences of *F. religiosa*. The parameters were set to identify perfect di-, tri-, tetra-, penta-, and hexa nucleotide motifs with a minimum threshold of 6, 5, 5, 5, and 5 repeats, respectively.

### Prediction of transcription factor families

The families of transcription factors (TFs) were predicted in genome annotations and differentially expressed transcripts of the Peepal tree using Plant Transcription Factor Database v5.0 [20].

### Non-coding RNA genes

The transfer RNAs in the Peepal tree genome were found using tRNAscan-SE (v2.0.3) [58] with the ‘eukaryotes’ option. tRNAscan-SE software deployed with the covariance models identifies the primary sequence and secondary structure information of tRNA and gives the complete tRNA genes for the query genome and transcriptome sequences. tRNAscan-SE software is integrated with Infernal v1.1 to enhance the tRNA search with better covariance and other updated models. Using the isotype-specific covariance model provides the functional classification of tRNAs and in the first pass search cutoff score, 10 is set. The miRbase database (http://www.mirbase.org) was used for the identification of putative miRNAs in the genome and unique identified transcripts sequence data based on the homology search. The long non-coding RNAs (lncRNAs) were identified with the Coding Potential Calculator tools [59].

### Transcriptome sequencing, assembly, and annotation

High-quality stranded RNA sequencing (ssRNA-seq) reads were assembled into putative transcripts using Trinity v2.9.0 [60]. Assembled transcripts were passed through Transdecoder v5.02 [61] to predict the coding sequences. The transcripts were clustered to find the unigenes by removing the redundant transcripts using the tool CD-HIT-est v.0.0.1 [62] with a 95% sequence identity threshold. Transcripts assembled from Trinity and CD-HIT-v0.0.1 were used in downstream analyses for gene prediction. Unigenes were used to predict the putative genes using the NCBI non-redundant (nr) database using the BLASTX program and proteins were predicted from the Uniprot database using the BLASTP program. The Trinity assembled transcripts were annotated using Trinotate-V3.11. The raw reads were mapped to scaffold assembled genome using Cufflinks-v2.2.1 [63] and considered as reference assembly.

### Transcript quantification and differential gene expression analysis

The estimation of transcripts abundance was determined using RNA-Seq by Expectation-Maximization (RSEM) tool [64], which quantifies transcript level abundance from RNA-seq data. RSEM first generates and pre-processes a set of reference transcript sequences and then aligns reads to reference transcripts followed by an estimation of transcript abundances. Normalized transcripts obtained from the transcript quantification methods were used in the next step for the differential gene expression analysis. FPKM and Trimmed Mean of M-values (TMM) are calculated to understand the expression levels of genes in day and night samples of the Peepal tree. For further analysis, the gene expression was estimated using FPKM and TMM value minimum ≥1. The TMM value was used to cluster the genes according to their expression pattern using the edgeR package in the R tool. The parameters used in the differential expression analysis were a probability value P-value of 0.001 and a fold change value of log2. The expression value was also determined for assembled transcripts to verify the expression of genes predicted from gene models. The differentially expressed genes were annotated using BLAST2GO Annotation software [65].

### Pathway Analysis

The annotated genes from the assembled genome and the differentially expressed genes from the Peepal tree leaf tissues collected during the day (2 PM) and night (2 AM) were used for pathway analysis in the KAAS (KEGG Automatic Annotation Server) (KEGG) server [17] using the BBH (bi-directional best hit) method and the search against a default set of 40 eukaryotic organisms. It provided the list of pathways where the candidate genes were mapped based on the orthologous homology alignment.

## Abbreviations

BUSCO: Benchmarking Universal Single-Copy Orthologous
ESTs: Expressed Sequence Tags
PPR repeat: Pentatricopeptide repeat
FPKM: Fragments Per Kilobase of transcript per Million mapped reads
MSA: Multiple Sequence Alignment

## Statement

- **Experimental Plant details:** *Ficus religiosa* a cultivated plant was used in this study. The mature leaves were collected from this plant at Private property, Anuganalu village, Hassan District, India (13.0647° N, 76.0363° E) were kindly provided by Prof. Malali Gowda, DNA Life Foundation, Anuganalu village, Hassan District, India (13.0647° N, 76.0363° E).
- The formal identification of the plant was undertaken by Prof. Malali Gowda, who owns the place of plant location and is also a corresponding author of this study. Hence, we had his permission to collect the plant material. The specimen of this material has not been deposited in any publicly available herbarium.

## Declaration

### Ethics approval and consent to participate

The study is compiled with relevant institutional, national, and international guidelines and legislation.

### Consent for publication

Not Applicable (NA)

### Availability of data and materials

The raw sequence reads have been deposited under NCBI Sequence Read Archive (SRA) accession numbers SRR7244210 for Illumina sequenced *F. religiosa* WGS, SRR13827064 for MGISEQ sequenced *F. religiosa* WGS – (https://trace.ncbi.nlm.nih.gov/Traces/study1/?acc=PRJNA474013), SRR7343291 (https://trace.ncbi.nlm.nih.gov/Traces/study1/?acc=SRR7343291) for *F. religiosa* transcriptome). The Whole Genome Shotgun project has been deposited at DDBJ/ENA/GenBank under the accession JAFMPE000000000 (https://www.ncbi.nlm.nih.gov/nuccore/JAFMPE000000000.1/) and Transcriptome Shotgun Assembly project has been deposited at DDBJ/ENA/GenBank under the accession GJAV00000000 (https://www.ncbi.nlm.nih.gov/nuccore/GJAV00000000.1/).

Other published data used in the study: The protein sequences of *Arabidopsis thaliana* (https://ftp.ncbi.nlm.nih.gov/genomes/all/GCF/000/001/735/GCF_000001735.4_TAIR10.1/GCF_000001735.4_TAIR10.1_protein.faa.gz), *Morus notabilis* (https://ftp.ncbi.nlm.nih.gov/genomes/all/GCF/000/414/095/GCF_000414095.1_ASM41409v2/GCF_000414095.1_ASM41409v2_protein.faa.gz), *Prunus persica* (https://ftp.ncbi.nlm.nih.gov/genomes/all/GCF/000/346/465/GCF_000346465.2_Prunus_persica_NCBIv2/GCF_000346465.2_Prunus_persica_NCBIv2_protein.faa.gz), *Cannabis sativa* (https://ftp.ncbi.nlm.nih.gov/genomes/all/GCF/900/626/175/GCF_900626175.2_cs10/GCF_900626175.2_cs10_protein.faa.gz), *Ziziphus jujuba* (https://ftp.ncbi.nlm.nih.gov/genomes/all/GCF/020/796/205/GCF_020796205.1_ASM2079620v1/GCF_020796205.1_ASM2079620v1_protein.faa.gz).

## Author’s contribution

AKL performed the DNA and RNA isolation from leaf tissues, genome assembly and functional annotation, gene prediction, repeat prediction, orthologous gene clustering, Genome and Transcriptome pathway analysis, other bioinformatic analysis submitted WGS and RNA-seq data to NCBI, prepared the genomic and transcriptomic study tables and figures, other bioinformatic analysis and wrote the manuscript; MG, AKL, and AKP designed the genomic and transcriptomic experiments of *Ficus religiosa*; MG conceived and conceptualized the project; reviewed and edited the manuscript. AKP reviewed and edited the manuscript. All authors have read and reviewed the final draft of the manuscript.

## Acknowledgment

We would like to thank Tata Education and Development Trust fellowship (01/2018-2019) provided for Ashalatha K L for her Ph.D. program. We acknowledge the Next Generation Genomics Facility at the Centre for Cellular and Molecular Platforms (C-CAMP) and Bengaluru Genomics Center Pvt. Ltd for support in sequencing; Mr. Ravindra Raut for his help in collecting the leaf samples.

